# Neural Network Guided Calibration for Fast Virtual Twin Generation in Cardiovascular ODE Models

**DOI:** 10.64898/2026.05.05.722845

**Authors:** MT Cabeleira, S Ray, NC Ovenden, V Diaz-Zuccarini

## Abstract

Calibration of closed-loop lumped-parameter cardiovascular models remains a major bottleneck for scalable digital-twin generation because inverse estimation is ill-conditioned and typically requires computationally expensive iterative forward simulation. This study investigates whether a supervised neural network (NN) can provide a fast inverse estimator for a paediatric sepsis cardiovascular ODE model by learning a direct mapping from prescribed haemodynamic target vectors to calibrated parameter sets.

Training data are generated by sampling model parameters at random, forward-simulating the closed-loop system to steady state, and pairing the resulting target summaries with the corresponding parameters; the same target definitions and evaluation populations are used throughout for consistency. We evaluate NN inference by forward re-simulation to steady state and benchmark performance against a simulator-constrained calibration reference (Embedded Gradient Descent, EGD) using relative-error statistics, distributional similarity of achieved outputs and inferred parameters (median shift, IQR ratio, Wasserstein distance, KS statistic), and target-space localisation of parameter-space disparity (cosine distance). The NN reproduces the prescribed targets with predominantly small errors for most samples, while the largest discrepancies are confined to a well defined set of target configurations that also yield high residuals under the reference method, indicating feasibility limits of the target/model combination.

Overall, NN-guided calibration provides a computationally efficient accelerator for virtual-twin generation and target-space screening, with simulator-based refinement and forward re-simulation retained to handle infeasible regimes and enforce mechanistic plausibility.

## 1 INTRODUCTION

Digital patient twins, computational models capable of reproducing an individual’s physiological dynamics, have emerged as a central paradigm in precision and patient-specific medicine. In particular, cardiovascular digital twins have developed rapidly, with recent publications [1, 2] outlining their system architectures, data-integration pipelines, and potential clinical applications within cardiovascular medicine. These efforts typically adopt either mechanistic models or machine-learning approaches to represent cardiovascular dynamics. Rather than treating these as competing paradigms, Holmes et al. [3] argue that they are fundamentally complementary, with mechanistic models providing causal structure, interpretability, and physiological constraints, whereas machine-learning methods offer computational efficiency, flexibility, and the ability to approximate high-dimensional non-linear relationships.

This complementarity is reflected across multiple domains of cardiovascular modelling. In cardiac electrophysiology and electromechanics, latent neural ODEs have been used to emulate the temporal dynamics of high-dimensional electromechanical models in a reduced latent space [4], and surrogate neural networks have been trained to approximate the outputs of full biophysical electromechanics solvers for real-time prediction [5]. In haemodynamics, emulator-based statistical models have been constructed on top of reduced-order mechanistic flow simulations to enable rapid exploration of parameter spaces [6]. Related efforts include hybrid neural-ODE approaches in which a lumped-parameter cardiovascular model is retained as the mechanistic backbone while only a specific non-linear interaction term is learned by a neural component [7]. Despite substantial progress in cardiovascular digital-twin modelling, the calibration of mechanistic cardiovascular models to patient-specific data remains a major outstanding challenge and a key limitation for scalable digital-twin deployment. Parameter estimation for lumped and closed-loop cardiovascular models has been shown to suffer from ill-conditioning, parameter correlations, and limited identifiability, even under idealised or synthetic measurement conditions [8, 9, 10].

The literature provides several strategies to calibrate these models to clinical data, including Bayesian and MCMC-based parameter-estimation pipelines [11], data-assimilation approaches that integrate model dynamics with sequential measurement updates [12], longitudinal calibration studies examining the stability and variability of inferred parameters when the model is repeatedly fitted to sequential haemodynamic data [13], and optimisation frameworks that combine sensitivity analysis with patient-specific parameter fitting [14]. Complementing these generic strategies, recent work has proposed calibration frameworks that exploit cardiovascular model structure through physiologically informed control mechanisms, enabling the simultaneous estimation of larger parameter sets than typically considered [15, 16]. A common observation across these approaches is the inherent difficulty of recovering physiologically coherent parameter sets from sparse haemodynamic measurements and the substantial computational demands associated with the calibration workflows. Given the considerable computational burden associated with these calibration methods, several studies have explored whether neural networks can serve as faster approximations to the inverse problem in cardiovascular modelling. Bonnemain et al. demonstrated that a supervised deep neural network can infer ventricular contractility parameters directly from arterial pressure waveforms [17]. Hanna et al. showed that neural-network surrogates of 0D cardiovascular models can accelerate parameter estimation and uncertainty quantification in workflows requiring repeated forward evaluations [18]. Physics-informed neural networks have also been used to solve both forward and inverse problems in physiological systems by embedding mechanistic constraints into the learning process [19], with applications such as cuffless blood-pressure reconstruction [20].

In this work, we investigate whether a neural-network-based inverse estimator can provide a fast and scalable alternative to optimisation-based mechanistic calibration for cardiovascular digital twin generation. Using the same population targets defined in our previous EGD study [16], we train a supervised neural network to approximate the inverse mapping from model outputs to the underlying parameter set. We then perform a systematic comparison between the neural-network estimator and the EGD framework, contrasting their behaviour across accuracy, computational cost, and physiological coherence. Unlike previous calibration studies, which typically estimate fewer than ten parameters simultaneously, both approaches in this work operate on a substantially larger parameter space. By examining the trade-off between the speed of the neural-network estimator and the precision of the mechanistic EGD method, we provide the first benchmark of data-driven and mechanistic inverse modelling approaches for generating coherent populations of cardiovascular digital twins.

## 2 MATERIALS AND METHODS

This section defines the inverse-calibration task posed by a closed-loop lumped-parameter cardiovascular model and the synthetic haemodynamic target populations used for evaluation. We first summarise the model and target definitions and outline the Embedded Gradient Descent (EGD) framework used as the mechanistic reference. We then describe the proposed neural-network inverse estimator, including dataset generation, architecture, training, and the inference plus forward re-simulation workflow. Finally, we specify the quantitative metrics used to compare methods, covering calibration error, distributional equivalence, and parameter-space disparity.

### 2.1 Calibration Problem Definition and Experimental Setup

#### 2.1.1 Cardiovascular Model Specification

The calibration problem considered in this work is defined on a closed-loop, lumped-parameter model of the cardiovascular system, comprising the heart and the systemic and pulmonary circulations. A schematic representation of the model structure is shown in Figure 1. The model represents the left and right heart chambers (Hl, Hr), systemic and pulmonary arteries (As, Ap), capillaries (Cs, Cp), veins (Vs, Vp), and thoracic veins (Vt) as interconnected compartments, each characterised by a pressure (*P*), a volume (*V*), and inflow and outflow rates 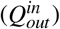 governed by resistive elements (*R*). The model operates as an electrical analogue, where compliant elements (*C*) represent vascular and cardiac capacitance and resistive elements govern blood transport between compartments. Time-varying elastance (*ℰ*) formulations are used to represent the contractile behaviour of the heart chambers, enabling cycle-resolved (*P*), (*V*), and flow (*Q*) dynamics to be generated throughout the closed-loop system.

**Figure 1.**
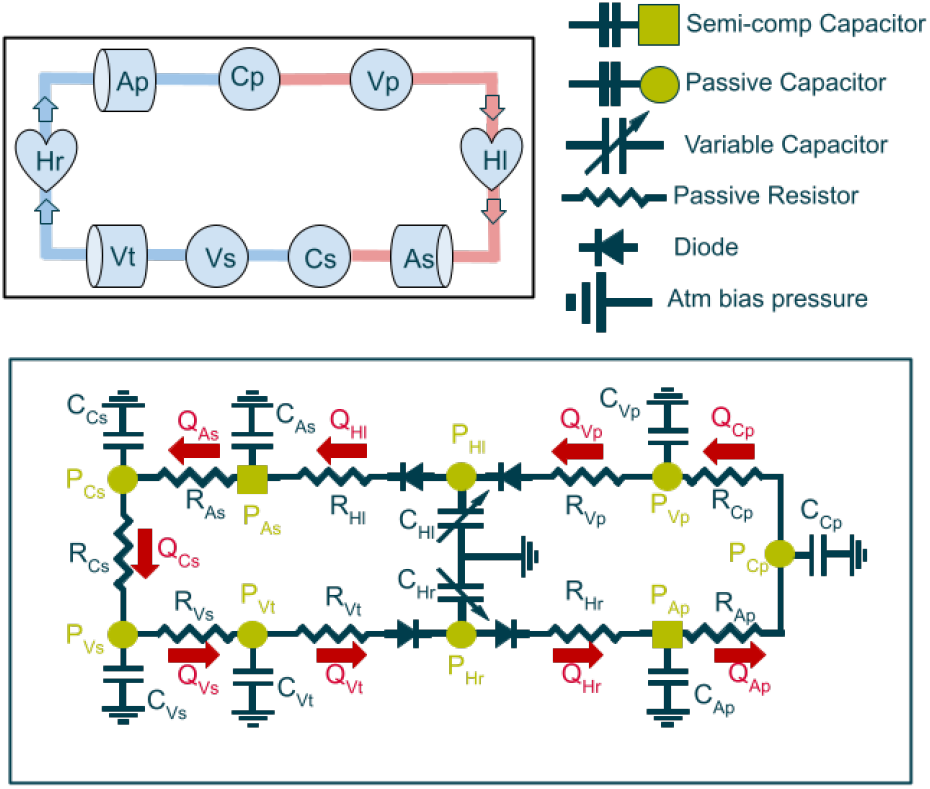
Schematic representation of the cardiovascular model used in this study. **Top:** High-level block diagram illustrating the closed-loop structure of the systemic and pulmonary circulations, including the left and right heart chambers (Hl, Hr), arterial (As, Ap), capillary (Cs, Cp), venous (Vs, Vp), and thoracic venous (Vt) compartments. **Bottom:** Detailed circuit representation of the same model, where each compartment is modelled as a compliant element and interconnected through resistive links that govern inter-compartmental blood flow. This representation defines the pressure/volume/flow relationships underlying the forward simulations used for calibration. A concise specification of the governing equations and cycle-based operators is provided in Appendix C.

The cardiovascular model used here is identical to that employed in our previous work [16]. A concise specification of the governing equations, cycle timing, and auxiliary operators used to compute calibration targets is provided in Appendix C. Full derivation, physiological justification, and validation of the model are reported in [16]. In the present study, the model is treated as a fixed forward simulator that defines the inverse calibration problem addressed by both the EGD and the current, neural-network-based, calibration methods.

#### 2.1.2 Synthetic Population and Calibration Targets

To compare calibration strategies on a realistic and clinically relevant problem, a synthetic population was generated for paediatric sepsis. This is a useful setting because septic shock is driven by interacting changes in vascular tone, cardiac function and total blood volume, whilst important internal quantities such as blood volume distribution, regional pressures and microcirculatory states cannot be measured directly from routine intensive care data. This is especially relevant in paediatric intensive care, where monitoring usually provides only a limited set of bedside variables and richer waveform data are not always available for routine use. Paediatric sepsis is also a particularly demanding calibration setting because it is heterogeneous, age-dependent and less well characterised than adult disease. It also includes distinct warm and cold shock phenotypes with different underlying haemodynamics but partly overlapping clinical signatures, making it both clinically important and well suited for testing whether a calibration method can recover plausible internal cardiovascular states from limited observable data.

The calibration targets used in this work are summarised in Table 1 and define the three benchmark states considered here. Target ranges were derived from the clinical literature and follow the same rationale introduced in our previous Embedded Gradient Descent (EGD) study [16], ensuring direct comparability between methods. In brief, the targets include cardiac output, heart rate, systemic and pulmonary arterial pressures (systolic and pulse), central venous pressure, pulmonary capillary pressure, and total blood volume.

**Table 1.**
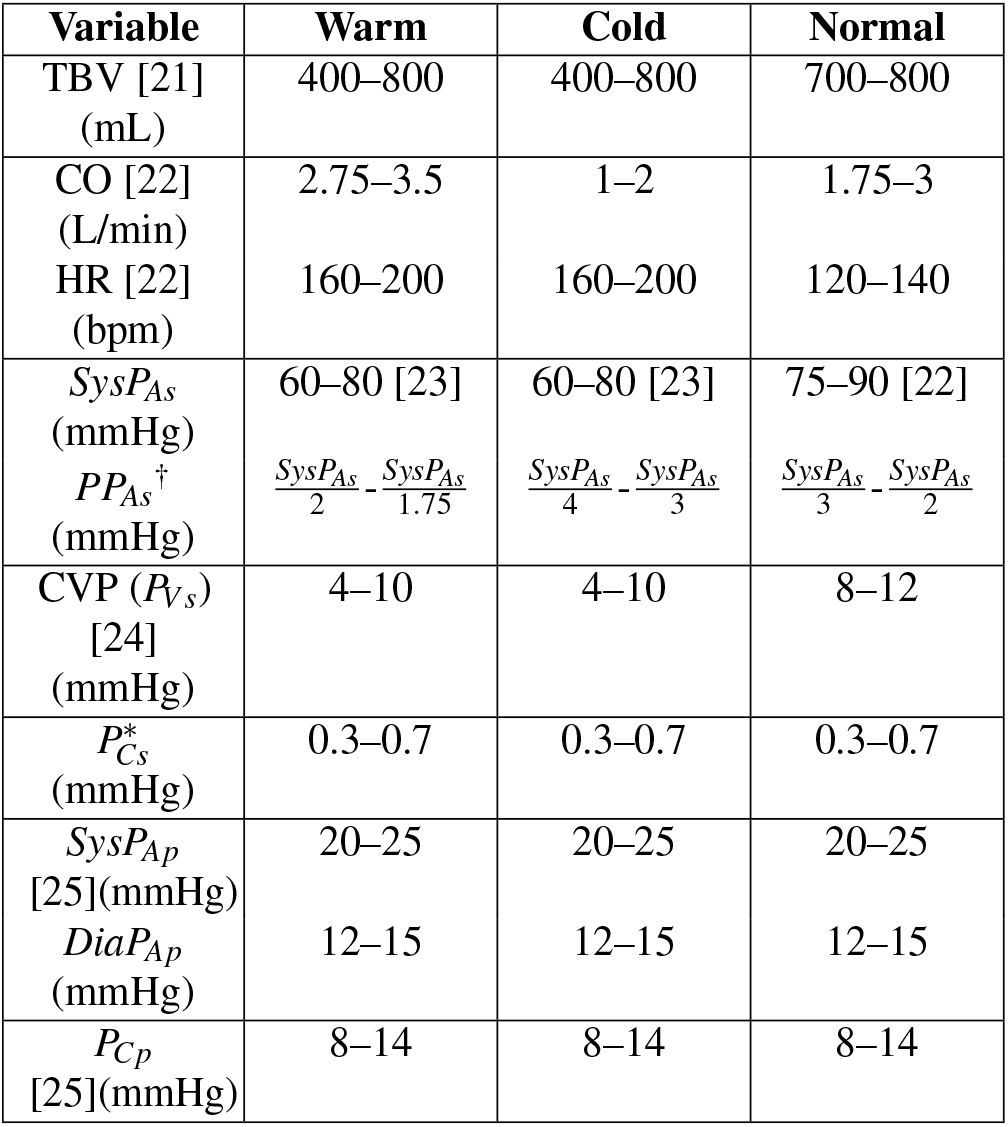
Target ranges used for calibration under warm shock, cold shock, and normal physiology. The compartment labels and symbols used in this table are illustrated in Figure 1 and listed in the nomenclature in Appendix B (Table 7).. As the capillary pressure of the systemic system is impossible to measure, we have imposed a % drop in pressure from the diastolic value of the Systemic arterial pressure and the value of CVP. † The targets for the Pulse Pressure of the systemic arterial blood pressure is calculated according to the equation extracted from Ranjit et al. [23] where these ratios were found to be descriptive of sepsis cases.

| Variable | Warm | Cold | Normal |
| --- | --- | --- | --- |
| TBV [21]<br>(mL) | 400–800 | 400–800 | 700–800 |
| CO [22]<br>(L/min) | 2.75–3.5 | 1–2 | 1.75–3 |
| HR [22]<br>(bpm) | 160–200 | 160–200 | 120–140 |
| $SysP_{As}$<br>(mmHg) | 60–80 [23] | 60–80 [23] | 75–90 [22] |
| $PP_{As}^{\dagger}$<br>(mmHg) | $\frac{SysP_{As}}{2} - \frac{SysP_{As}}{1.75}$ | $\frac{SysP_{As}}{4} - \frac{SysP_{As}}{3}$ | $\frac{SysP_{As}}{3} - \frac{SysP_{As}}{2}$ |
| CVP ($P_{Vs}$)<br>[24]<br>(mmHg) | 4–10 | 4–10 | 8–12 |
| $P_{Cs}^*$<br>(mmHg) | 0.3–0.7 | 0.3–0.7 | 0.3–0.7 |
| $SysP_{Ap}$<br>[25](mmHg) | 20–25 | 20–25 | 20–25 |
| $DiaP_{Ap}$<br>(mmHg) | 12–15 | 12–15 | 12–15 |
| $P_{Cp}$<br>[25](mmHg) | 8–14 | 8–14 | 8–14 |

In addition to pressure and flow-based targets, the calibration procedure relies on assumptions regarding blood volume distribution across the cardiovascular compartments. Average compartmental volume fractions, reported in Table 2, were adopted as physiological priors and scaled according to the prescribed total blood volume (TBV). This fixed distribution provides a consistent reference across the synthetic population and avoids introducing additional degrees of freedom during calibration. As discussed extensively in [16], this choice facilitates population-level comparison but may restrict compatibility with certain target combinations under extreme haemodynamic conditions.

**Table 2.** Average blood volumes on different parts of the circulatory system, assuming a total blood volume of 700mL. Extracted from [21].

| Cardiovascular Region | Volumes |  |
| --- | --- | --- |
|  | mL | % |
| Left Heart | 24.5 | 3.5 |
| Right Heart | 24.5 | 3.5 |
| Systemic arteries (Aorta) | 91 | 13 |
| Systemic arterioles & capillaries | 49 | 7 |
| Systemic Veins | 448 | 64 |
| Pulmonary Arteries | 26.6 | 3.8 |
| Pulmonary Capillaries | 9.8 | 1.4 |
| Pulmonary Veins | 26.6 | 3.8 |

Particular care was taken in defining targets for capillary pressures, which are not directly measurable in clinical practice. Systemic capillary pressure targets were, therefore, specified indirectly as a fractional pressure drop between diastolic systemic arterial pressure and central venous pressure.

The target definitions represent physiologically plausible scenarios but remain approximate and may contain internal inconsistencies arising from simplified or population-level assumptions. In this study, the same target set is retained without modification in order to ensure a direct and controlled comparison between the reference Embedded Gradient Descent calibration and the proposed neural-network-based inverse calibration method.

#### 2.1.3 Reference Calibration Method: Embedded Gradient Descent

As a reference calibration approach, we employ the Embedded Gradient Descent (EGD) framework introduced in our previous work [16]. In that study, EGD was developed as a physiologically-informed optimisation strategy capable of calibrating a closed-loop cardiovascular model to a broad set of haemodynamic targets simultaneously.

EGD formulates calibration as a dynamic feedback process embedded directly within the model equations. Calibration variables are adjusted online through controller-like update laws that act on physiologically meaningful parameter groups, driven by the discrepancy between simulated outputs and prescribed target ranges. Each controller is associated with a specific physiological target category and adjusts a predefined subset of model parameters with a fixed correlation sign, as summarised in Table 3.

**Table 3.** Physiological calibration targets, corresponding observables, and dominant model parameters adjusted during calibration. The correlation sign denotes the qualitative parameter/observable relationship used in the controller de-sign: + for a positive correlation and − for a negative correlation.Overbars denote cycle-averaged quantities.

| Physiological target category | Calibration observables ($\mathbf{Y}$) | Primary parameters adjusted ($\hat{\theta}$) | Corr. |
| --- | --- | --- | --- |
| Heart volumes | $\bar{V}_{HL}, \bar{V}_{Hr}$ | $e_{HL}, e_{Hr}$ | - |
| Arterial volumes | $\bar{V}_{As}, \bar{V}_{Ap}$ | $V_{0As}, V_{0Ap}$ | + |
| Venous and capillary volumes | $\bar{V}_{Vt}, \bar{V}_{Cs}, \bar{V}_{Cp}, \bar{V}_{Vp}$ | $C_{Vt}, C_{Cs}, C_{Cp}, C_{Vp}$ | + |
| Cardiac output | $SV$ | $E_{Hl}$ | + |
| Systolic systemic arterial pressure | $SysP_{As}$ | $R_{AsCs}$ | + |
| Systolic pulmonary arterial pressure | $SysP_{Ap}$ | $E_{Hr}$ | + |
| Pulse pressures | $PP_{As}, PP_{Ap}$ | $C_{As}, C_{Ap}$ | - |
| Central venous pressure | $\bar{P}_{Vs}$ | $C_{Vs}$ | - |
| Systemic capillary pressure | $\bar{P}_{Cs}$ | $R_{CsVs}$ | + |
| Pulmonary capillary pressure | $\bar{P}_{Cp}$ | $R_{ApCp}$ | - |

In the present work, the EGD framework is used without modification. The same set of calibration observables, target definitions, and parameter groupings are retained, ensuring that both EGD and the proposed neural-network-based approach solve an identical inverse problem. Full details of the controller definitions, update equations, and numerical implementation are reported in [16] and are not repeated here.

The EGD study demonstrated that this approach can successfully recover physiologically coherent parameter sets across normal, warm shock, and cold shock regimes, even when calibrating a substantially larger number of parameters than typically considered in cardiovascular inverse problems. However, it also highlighted two key limitations relevant to the present comparison: the high computational cost associated with iterative forward simulation, and the presence of persistent residual errors for certain target categories arising from structural incompatibilities between imposed targets and fixed modelling assumptions.

### 2.2 Neural Network Inverse Calibration Method

To address the computational cost associated with repeated forward simulations in the EGD framework, we introduce a neural-network-based inverse calibration method. The proposed approach is designed to approximate the steady-state parameter mappings produced by EGD, replacing the iterative, controller-driven calibration process with a direct regression from physiological target space to model parameter space.

#### 2.2.1 Inverse problem formulation

Let **Y** ∈ ℝ^*M*^ denote the vector of physiological target quantities provided as input to the neural-network-based inverse calibration method, where *M* is the number of target quantities. These targets correspond to the haemodynamic observables listed in Table 1, augmented by total blood volume (TBV) together with its prescribed distribution across cardiovascular compartments (Table 2). All target quantities are defined as cycle-averaged or cycle-derived measures computed from forward simulations of the cardiovascular model.

In the Embedded Gradient Descent (EGD) framework, calibration is driven by a subset of these observables, resulting in an effective target space 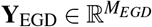, where *M*_*EGD*_ = 16, as defined by the controller-target mapping in Table 3. Two additional quantities are treated implicitly in EGD through initial conditions and fixed constraints. In particular, heart rate is prescribed as an initial condition for the forward simulation, and the average systemic venous volume 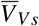 is indirectly constrained via the imposed total blood volume and compartmental volume distribution.

In contrast, the neural-network-based approach does not operate within a forward simulation during inference and therefore cannot rely on implicit initial conditions. To preserve equivalence with the EGD calibration task, heart rate (*HR*) and the systemic venous volume 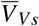 are explicitly included in the neural network input, resulting in an *M*_*NN*_-dimensional target vector, where *M*_*NN*_ = 18. The output of both calibration approaches is identical, consisting of the same 16-dimensional parameter vector θ ∈ ℝ^16^ defined and labelled by Table 3.

The neural network is trained to learn a mapping

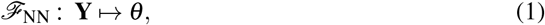

which approximates the steady-state parameter configurations obtained under the EGD calibration dynamics.

#### 2.2.2 Training dataset construction

The training dataset for the neural-network inverse calibration method was constructed to mirror the calibration task addressed by the Embedded Gradient Descent (EGD) framework, while making all target quantities explicit. In contrast to EGD, where part of the calibration problem is handled implicitly through initial conditions and fixed constraints, the neural network operates on an explicit target vector **Y** ∈ ℝ^18^, as defined in Section 2.2.1. All target quantities used for training therefore correspond directly to the inputs provided to the network at inference time.

Parameter ranges used for data generation were derived from the calibrated populations obtained using the EGD method. For each calibrated parameter, the minimum and maximum values observed across the EGD population were extracted and subsequently expanded to define broader sampling bounds. This expansion was introduced to improve coverage of the admissible parameter space and to reduce boundary effects during network training. The resulting lower and upper bounds for each parameter are reported in Table 4. Using these bounds, 240,000 parameter vectors were generated via Latin hypercube sampling to ensure uniform coverage of the high-dimensional parameter space. Each sampled parameter set was run through the cardiovascular model until a periodic steady state was reached (20 seconds). From each simulation, the corresponding physiological target vector **Y** were then extracted. These correspond to the values of the variables in Table 3 at the end of each simulation, once periodic steady state had been reached.

**Table 4.** Observed parameter ranges in the Sepsis population and corresponding bounds used for the generation of the training set. *HC* and *TBV* are also used here to define the initial heart-rate and total blood volume in the system.

| Param ($\hat{\theta}$) | $Min_{EGD}$ | $Max_{EGD}$ | $LowB_{NN}$ | $UppB_{NN}$ |
| --- | --- | --- | --- | --- |
| $C_{Vs}$ | 25.04 | 115.88 | 10.0 | 350 |
| $R_{ApCp}$ | 0.08 | 0.54 | 0.03 | 2.0 |
| $R_{CsVs}$ | 0.11 | 1.75 | 0.02 | 2.5 |
| $R_{AsCs}$ | 0.41 | 2.58 | 0.10 | 4.5 |
| $E_{HI}$ | 3.44 | 30.00 | 0.5 | 50.0 |
| $E_{Hr}$ | 1.00 | 9.09 | 0.3 | 20.0 |
| $C_{As}$ | 0.16 | 0.52 | 0.05 | 1.8 |
| $C_{Ap}$ | 0.22 | 2.43 | 0.05 | 4.0 |
| $V_{OAs}$ | 35.01 | 92.00 | 10.0 | 200 |
| $V_{OAp}$ | 1.00 | 25.52 | 0.0 | 40.0 |
| $e_{HI}$ | 0.22 | 0.95 | 0.05 | 1.8 |
| $e_{Hr}$ | 0.06 | 0.66 | 0.02 | 1.8 |
| $C_{Cs}$ | 0.76 | 4.43 | 0.3 | 10.0 |
| $C_{Vt}$ | 1.94 | 12.68 | 0.5 | 25.0 |
| $C_{Cp}$ | 0.75 | 2.53 | 0.3 | 6.0 |
| $C_{Vp}$ | 1.09 | 3.74 | 0.5 | 8.0 |
| $HC$ | 0.30 | 0.50 | 0.25 | 0.60 |
| $TBV$ | 400 | 700 | 350 | 1000 |

Simulations that failed to converge or produced non-physiological outputs were discarded during a preprocessing stage. The remaining samples formed paired training data of the form (**Y**, *θ*), where *θ* denotes the underlying parameter vector used in the forward simulation.

The processed dataset was subsequently split into training, validation, and test subsets using an 80/15/5 partition. Both input targets and output parameters were standardised using zero-mean, unit-variance normalisation, with scaling statistics computed exclusively from the training subset and subsequently applied unchanged to the validation and test sets.

#### 2.2.3 Network architecture

The inverse calibration mapping *ℱ* _NN_: **Y** → *θ* was implemented as a fully connected feedforward neural network using TensorFlow/Keras. The network takes as input the 18-dimensional target vector **Y** ∈ ℝ^18^ (Section 2.2.1) and outputs a 16-dimensional parameter vector 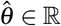^16^, corresponding to the calibrated parameter set defined in Table 3.

##### Layer configuration

The architecture comprises 3 hidden layers of equal width (128 units each), followed by a linear output layer:

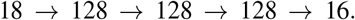

All hidden layers use a hyperbolic tangent activation function, which provides bounded, symmetric nonlinearities well suited to the zero-mean, unit-variance normalised input and output spaces used during training. This choice stabilises optimisation for the ill-conditioned inverse calibration problem while allowing the output layer, which uses a linear activation, to represent unbounded parameter values after de-standardisation.

##### Objective function and optimization

Training was performed using a mean-squared-error loss defined in parameter space,

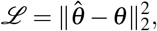

where both 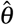 and *θ* are expressed in the standardised coordinate system defined by the output scaler (zero mean and unit variance, fitted on the training set only; Section 2.2.2). A mean-squared-error loss in parameter space was used, reflecting the objective of approximating the calibrated parameter values produced by the Embedded Gradient Descent framework rather than directly minimising output-level discrepancies. This choice ensures balanced contributions from all calibrated parameters and yields a smooth optimisation landscape for the inverse regression task. The optimisation used the Adam optimiser [26] with an exponentially decaying learning rate schedule. The initial learning rate was set to 10^−3^ and decayed by a factor of 0.9 every 60,000 optimisation steps using a staircase schedule. These optimisation hyperparameters were selected empirically based on validation performance and training stability.

##### Training protocol

Models were trained with a batch size of 256 for up to 1000 epochs. Early stopping was applied based on the validation loss, with a patience of 50 epochs and a minimum improvement threshold of 5 × 10^−5^. The weights corresponding to the best validation loss were restored at the end of training.

#### 2.2.4 Inference and forward re-simulation

During inference, prescribed physiological target vectors **Y**_target_ are first standardised using the scaling parameters estimated from the training dataset and then passed through the trained neural network to obtain predicted parameter vectors 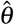. These predictions are subsequently de-standardised and interpreted as calibrated model parameters.

To evaluate the quality and physiological consistency of the inferred parameters, each predicted parameter set is reinjected into the cardiovascular model and propagated forward in time until a steady-state solution is reached. The resulting haemodynamic outputs are then compared against the prescribed target quantities. This forward re-simulation step does not constitute an additional calibration stage, but rather serves as a validation procedure to assess inverse accuracy, generalisation, and consistency with the underlying mechanistic model.

This inference and validation workflow corresponds to Steps 2 and 3 in Figure 2. Because the neural network operates directly in target space, the same procedure can be applied to arbitrary admissible target sets, enabling rapid generation and evaluation of calibrated parameter populations without iterative optimisation.

**Figure 2.**
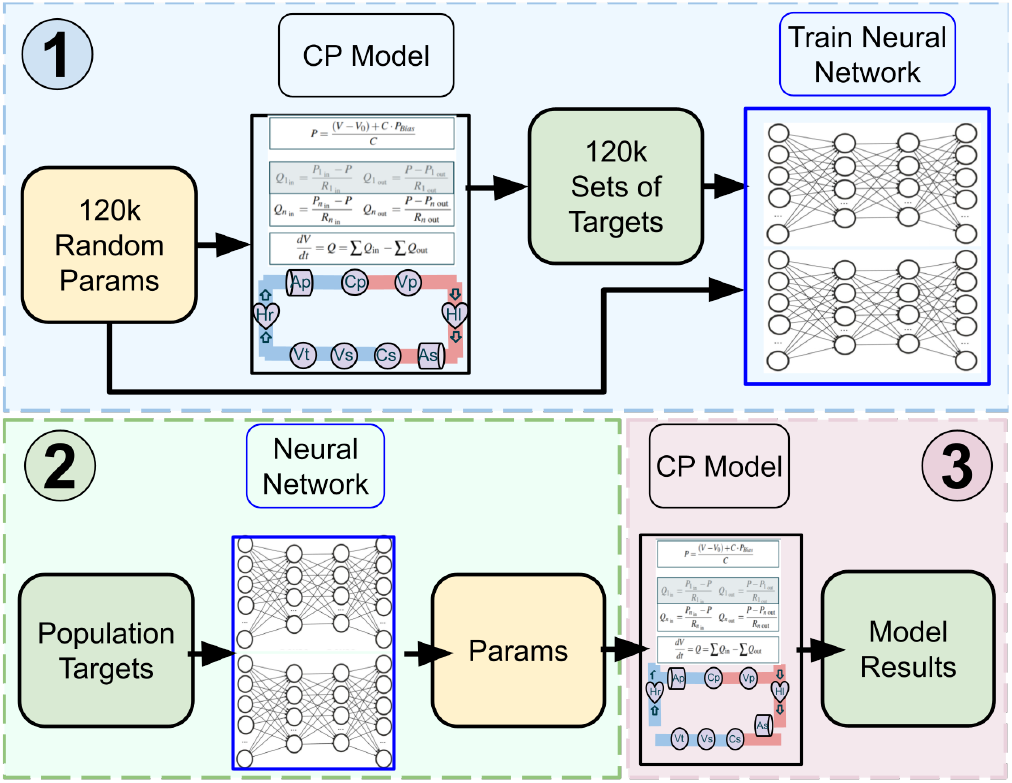
Overview of the neural-network-based inverse calibration workflow. (1) Model parameters are sampled within physiological bounds using Latin hypercube sampling and propagated through the CP model to generate corresponding haemodynamic target quantities. These target/parameter pairs are used to train a supervised neural network that learns an approximate inverse mapping from physiological targets to model parameters. (2) Prescribed physiological target sets are provided as inputs to the trained neural network, which directly infers the corresponding calibrated model parameters in a single forward pass, without requiring iterative forward simulations. (3) The inferred parameter sets are re-injected into the CP model to generate forward simulations. The resulting haemodynamic outputs are compared against the prescribed targets to quantify inverse accuracy, assess generalisation, and evaluate physiological consistency relative to the reference EGD calibration.

### 2.3 Calibration Comparison Methods

To enable a controlled and reproducible comparison between the Embedded Gradient Descent (EGD) framework and the proposed neural-network (NN) inverse calibration method, all simulations were performed using the same calibration targets, model structure, and evaluation protocol.

#### Target generation and population sampling

Calibration targets were generated using the same physiological ranges defined in the EGD study. For each phenotype (normal physiology, warm septic shock, and cold septic shock), Latin hypercube sampling (LHS) was used to uniformly sample the admissible target space within the prescribed bounds of Table 1. This approach ensures uniform coverage of the multidimensional haemodynamic target domain. For each phenotype, 200 independent target sets were generated. Because the same target sets were used for both the EGD and NN calibration runs, any discrepancies observed between methods arise solely from differences in the calibration strategy, not from differences in target definition, sampling, or model configuration.

#### EGD calibration runs

For each sampled target vector, the EGD framework was initialised with randomised parameter values within physiological bounds. The controller equations described in [16] were embedded directly within the cardiovascular model ODE system. Each controller adjusts a predefined parameter group according to the signed relative error between simulated and target observables. Simulations were propagated forward in time until convergence to a steady state was achieved. Convergence was defined operationally as stabilisation of cycle-averaged observables and saturation of controller dynamics within prescribed parameter bounds. The resulting steady-state parameter vector and haemodynamic outputs were stored for analysis.

#### Neural-network inference runs

The identical target sets generated via LHS were then provided as inputs to the trained neural network. Unlike EGD, the NN produces a calibrated parameter vector in a single forward pass without iterative simulation. To quantify calibration accuracy, each NN-predicted parameter set was re-injected into the cardiovascular model and propagated forward to steady state. This forward simulation step allows direct computation of calibration error by comparing the resulting haemodynamic outputs against the prescribed target values.

#### Calibration error summary statistics

For each calibrated sample and each physiological observable *Y*, relative error (*σ*) was computed after forward re-simulation as

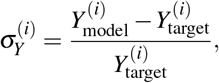

where *i* indexes population samples. From the distribution of 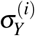 across samples, the following summary statistics were computed: Mean Absolute Error (MAE_*Y*_ = 𝔼 [❘ *σ*_*Y*_ ❘]) Standard Deviation (SD_*Y*_) and Root Mean Square Error 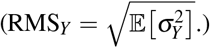.

Relative error was used to ensure scale invariance across observables with different physical units and magnitudes, allowing calibration performance to be compared directly across heterogeneous physiological targets. MAE quantifies the typical magnitude of deviation irrespective of sign, RMS emphasises larger deviations and is therefore sensitive to occasional calibration failures, and SD characterises the dispersion of signed errors across the virtual population. For a well-converged and unbiased calibration, the mean of *σ*_*Y*_ should be approximately zero, MAE_*Y*_ and RMS_*Y*_ should be small (e.g., *<*1–5% for tight physiological agreement), and RMS_*Y*_ should not substantially exceed MAE_*Y*_, which would otherwise indicate heavy-tailed residuals or unstable convergence in a subset of samples.

#### Distributional equivalence analysis

To assess whether the NN reproduces the population-level behaviour generated by EGD, we compared the empirical distributions of steady-state variables and calibrated parameters across the virtual population. Similarity was quantified using complementary statistics capturing location bias, dispersion changes, and distributional shape.

Systematic bias was assessed using the relative median shift,

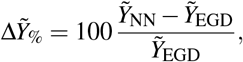

where 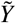 denotes the sample median; values near 0 indicate no population-level offset between methods. Preservation of variability was evaluated using the interquartile range (IQR) ratio,

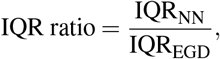

with values near 1 indicating matched dispersion (ratios *<* 1 indicate compression of variability and ratios *>* 1 indicate inflation).

Global distributional divergence in native units was quantified using the first-order Wasserstein distance,

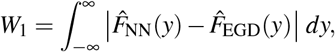

where 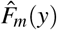 is the empirical cumulative distribution function for method *m*; *W*_1_ reflects the typical magnitude of population-level shift required to align the two distributions and is therefore interpretable in the units of *Y*. Differences in distribution shape were further assessed using the Kolmogorov–Smirnov test,

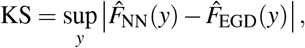

which measures the maximum vertical separation between the two empirical cumulative distribution functions and is bounded in [0, 1].

The two metrics capture distinct aspects of divergence. *W*_1_ aggregates differences across the full support of the distribution and is sensitive to broad population-level shifts, whereas KS detects the single largest local discrepancy, including deviations concentrated in specific regions (e.g. distribution tails). Using both ensures that a distribution is not considered preserved merely because average transport is small while a pronounced local structural deviation remains.

#### Parameter-space disparity analysis

Although forward haemodynamic agreement may be preserved, the NN may recover parameter vectors that differ from those obtained using EGD for identical targets. To quantify agreement in parameter-space structure between methods, we computed the cosine distance between the calibrated parameter vectors,

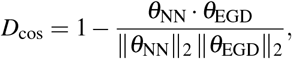

evaluated per calibrated sample using the full parameter vector.

Cosine distance measures the angular separation between parameter vectors and is therefore insensitive to global scaling, capturing whether the NN preserves the relative pattern of parameters (i.e., ratios/trade-offs across compartments) implied by the EGD solution. In this setting, low *D*_cos_ indicates near-collinearity and hence strong structural agreement in how parameters are configured to achieve the specified targets, whereas elevated *D*_cos_ indicates that the NN is producing a qualitatively different parameterisation for the same target configuration. Computing *D*_cos_ as a function of target configuration enables localisation of regions in target space where the inverse mapping is consistent between methods versus regions where alternative parameter patterns emerge; relating *D*_cos_ to forward haemodynamic error further distinguishes benign differences under accurate forward reproduction from discrepancies associated with degraded physiological fit.

## 3 RESULTS

The performance of the EGD and NN inverse calibration methods was first assessed by forward simulation on each calibrated sample. Table 5 reports, for each physiological observable, three population-level summary statistics of the post-calibration relative error for each method: mean absolute error (MAE), standard deviation (SD), and root mean square error (RMS), shown separately for EGD and NN and expressed as percentages. Here, we see consistently lower MAE, SD, and RMS for EGD across all reported observables, indicating both higher accuracy and more stable convergence, whereas the NN exhibits markedly larger errors for most pressures and especially for venous and peripheral volume observables (e.g., 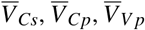). Similarly to the EGD method, the most challenging variable is 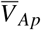, which shows the largest variability (SD ≈ 13–14%) and similarly poor RMS in both cases. We also quantify how closely the NN reproduces the EGD population distribution for each sample using complementary measures of location shift 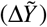, dispersion (IQR ratio), and global distributional discrepancy (*W*_1_, KS). Overall, the achieved observable distributions are closely matched; for most targets 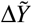 is near zero and the IQR ratio is close to unity, indicating minimal population-level bias and similar dispersion between methods. The largest discrepancies occur for venous/peripheral volume observables, particularly 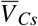 and 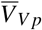, which show the strongest median shifts and the highest *W*_1_/KS values, consistent with the broader error behaviour observed for these compartments.

**Table 5.** Calibration error summary statistics for EGD and NN calibration. For each physiological observable, the table reports the mean absolute relative error (MAE), the standard deviation of relative error (SD), and the root mean square relative error (RMS) computed across the calibrated population after forward simulation. All values are expressed as percentages, enabling direct comparison of accuracy (MAE), precision (SD), and sensitivity to outliers (RMS) between methods. For each variable, the table also reports the relative median shift 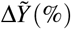, the interquartile range (IQR) ratio, the first-order Wasserstein distance *W*_1_, and the Kolmogorov–Smirnov (KS) statistic comparing the NN and EGD population distributions.

| Var. | EGD |  |  | NN |  |  | Comparison |  |  |  |
| --- | --- | --- | --- | --- | --- | --- | --- | --- | --- | --- |
| | MAE | SD | RMS | MAE | SD | RMS | $\Delta\tilde{Y}$ (%) | IQR ratio | $W_1$ | KS |
| $\bar{P}_{Vs}$ | 0.27 | 0.31 | 0.32 | 2.13 | 1.79 | 2.49 | +0.03 | 1.014 | 0.143 | 0.057 |
| $\bar{P}_{Cp}$ | 0.78 | 1.64 | 1.64 | 2.76 | 3.59 | 3.59 | -0.01 | 1.014 | 0.0859 | 0.033 |
| $\bar{P}_{Cs}$ | 0.70 | 1.16 | 1.17 | 3.30 | 3.98 | 4.08 | -0.01 | 0.995 | 0.262 | 0.050 |
| $P_{As}^{\text{max}}$ | 0.64 | 1.16 | 1.17 | 3.23 | 3.95 | 4.05 | -0.02 | 1.127 | 0.836 | 0.077 |
| $SV$ | 0.94 | 2.36 | 2.38 | 3.83 | 3.98 | 4.67 | +2.56 | 1.032 | 0.466 | 0.063 |
| $P_{Ap}^{\text{max}}$ | 0.70 | 1.11 | 1.14 | 2.91 | 3.75 | 3.79 | +0.01 | 1.107 | 0.291 | 0.102 |
| $PP_{As}$ | 0.22 | 0.87 | 0.87 | 3.96 | 5.07 | 5.12 | +1.59 | 1.055 | 0.609 | 0.062 |
| $P_{Ap}^{\text{min}}$ | 1.26 | 1.99 | 2.05 | 3.41 | 4.23 | 4.57 | -0.02 | 1.012 | 0.193 | 0.078 |
| $\bar{V}_{As}$ | 0.34 | 0.48 | 0.49 | 1.01 | 1.30 | 1.33 | +0.01 | 0.995 | 0.317 | 0.032 |
| $\bar{V}_{Ap}$ | 4.57 | 14.19 | 14.77 | 7.15 | 13.06 | 14.32 | +0.60 | 1.072 | 0.576 | 0.113 |
| $\bar{V}_{Hl}$ | 1.92 | 2.54 | 2.54 | 3.66 | 4.64 | 4.77 | +0.44 | 1.057 | 0.2 | 0.068 |
| $\bar{V}_{Hr}$ | 1.82 | 2.34 | 2.34 | 3.55 | 3.69 | 4.42 | +2.37 | 1.032 | 0.394 | 0.087 |
| $\bar{V}_{Cs}$ | 0.74 | 0.91 | 0.91 | 5.02 | 5.17 | 6.34 | -5.66 | 0.976 | 1.75 | 0.143 |
| $\bar{V}_{Vt}$ | 0.52 | 0.63 | 0.63 | 3.08 | 3.14 | 3.76 | +0.78 | 1.012 | 0.587 | 0.107 |
| $\bar{V}_{Cp}$ | 0.45 | 0.56 | 0.56 | 4.02 | 5.08 | 5.34 | -2.58 | 1.021 | 0.331 | 0.087 |
| $\bar{V}_{Vp}$ | 0.45 | 0.55 | 0.56 | 5.01 | 6.15 | 6.31 | -1.14 | 1.204 | 0.709 | 0.178 |
| $V_{Vs}$ | 0.53 | 1.04 | 1.08 | 0.98 | 1.23 | 1.27 | -0.20 | 0.962 | 1.33 | 0.037 |

Figure 3 summarises target/output agreement (scatter about the identity line) and the corresponding population distributions of relative error (violin plots) for a representative subset of calibration variables. In Appendix A, the full set of parameters are displayed. The subset is chosen to illustrate key regimes highlighted by Table 5: *P*_*Vs*_ as the best-tracked variable (lowest errors), 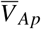 as the most challenging variable (largest dispersion and heavy-tailed residuals), *V*_*Hl*_ as a higher-error case where both methods remain broadly comparable, *V*_*Cs*_ because it exhibits one of the strongest median shifts, and *SV* as a central flow-derived target. Overall, EGD achieves tighter target tracking and more concentrated error distributions than NN across these observables, indicating improved accuracy and reduced dispersion across the population. Despite this, NN predictions form a dense cloud around the identity line for most targets, with the majority of samples exhibiting relatively small errors and relatively few extreme outliers. 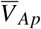 is the main exception: while most NN samples remain comparable to the other volume targets, a small subset exhibits very large deviations, producing a heavy-tailed error distribution; for this variable the NN shows fewer extreme outliers than EGD.

**Figure 3.**
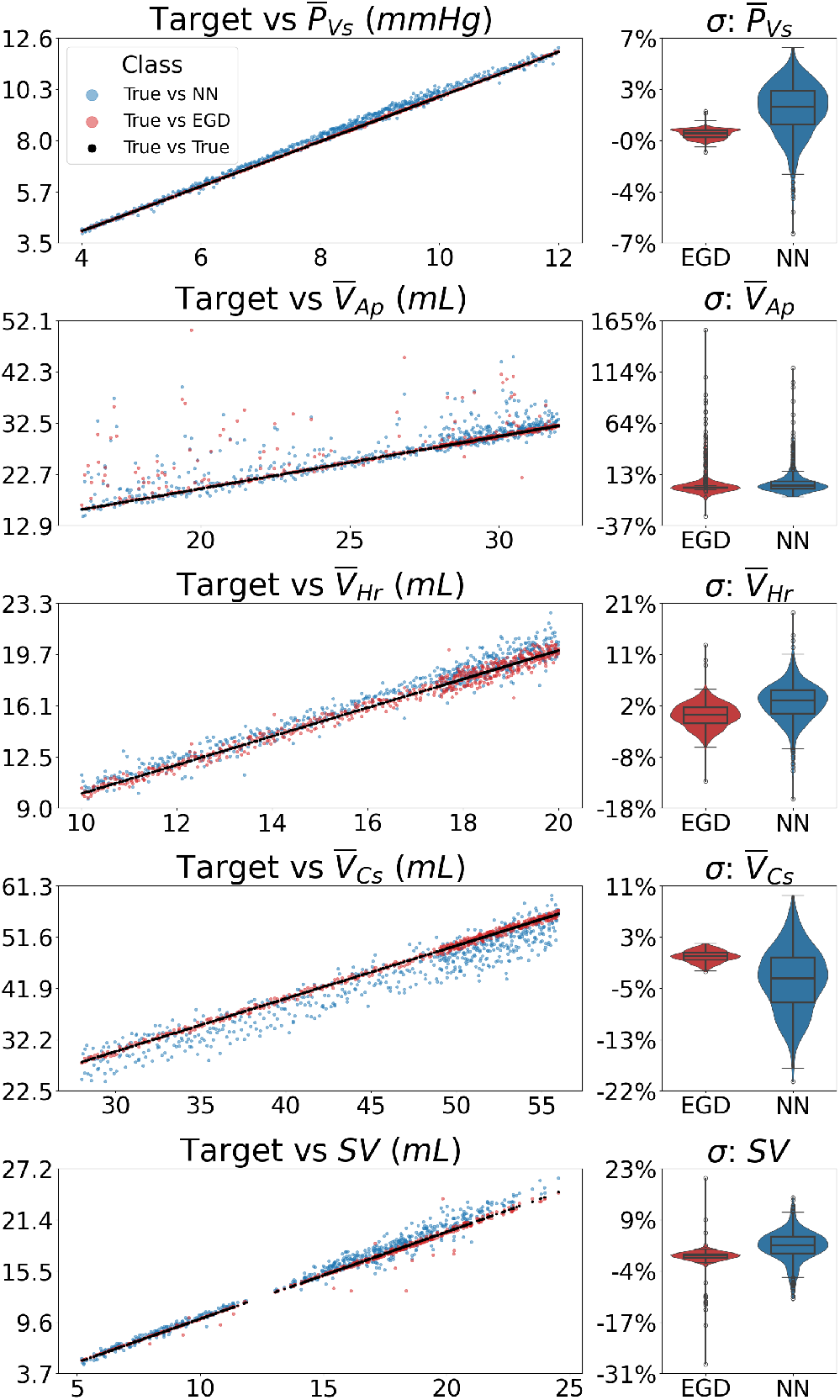
Target/output agreement and relative-error distributions for a representative subset of calibration variables. For each variable *Y*, the scatter plot compares prescribed targets against forward re-simulated steady-state outputs (identity line), while the adjacent violin plot shows the corresponding distribution of relative error *σ*_*Y*_ = (*Y*_model_ −*Y*_target_)*/Y*_target_ across the calibrated population for EGD and NN.

To investigate whether the differences in calibration performance arise from systematic differences in the inferred parameter distribution, we compare the population distributions of the calibrated model parameters obtained using EGD and the NN (Figure 4). For each calibrated parameter listed in Table 3, the population distribution obtained from EGD (red) and from the NN inverse mapping (blue), highlighting agreement or systematic shifts in location (median), dispersion (IQR), and tail behaviour (outliers). Across many parameters, the NN reproduces the central tendency of the EGD population well, with comparable medians and interquartile ranges, indicating that the inverse mapping preserves the bulk of the calibrated parameter space.

**Figure 4.**
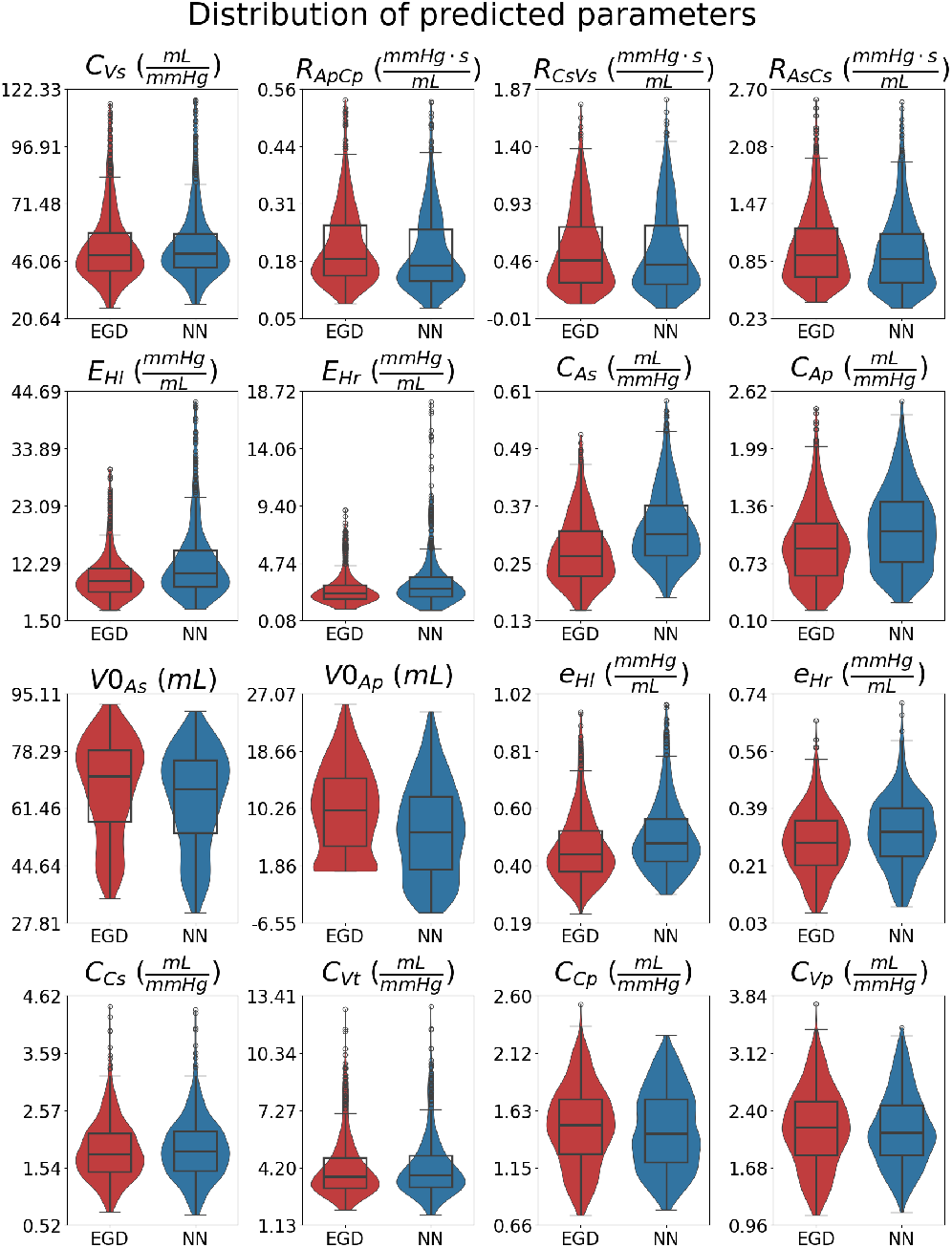
Population distributions of calibrated model parameters obtained using Embedded Gradient Descent (EGD) and neural-network (NN) inverse calibration. Each panel shows the empirical distribution across the virtual population for one parameter, with violin plots indicating density and overlaid box plots indicating the median and interquartile range.

However, systematic shifts are apparent for specific parameter groups: the NN tends to predict slightly lower values for the resistance parameters (*R*_*ApCp*_, *R*_*CsVs*_, *R*_*AsCs*_) and higher values for arterial compliance parameters (*C*_*As*_, *C*_*Ap*_), consistent with a structured rebalancing of vascular load between resistance and compliance. The most pronounced discrepancies occur in the heart elastance parameters (*E*_*Hl*_, *E*_*Hr*_), where the NN exhibits substantially broader distributions and heavier upper tails, suggesting occasional extrapolation to higher contractility states relative to EGD. Unstressed volume parameters, particularly *V*0_*Ap*_, also show a noticeable shift and increased spread. We can also see that the *V*0_*Ap*_ is shifted into negative values which would be inconsistent with physiology as this parameter represents the volume the vessel occupies when no pressure differential between the walls is applied.

Table 6 compares the population distributions of calibrated parameters inferred by NN and EGD, highlighting whether the NN recovers the same parameter-space structure (median, spread, and distributional shape) as the EGD-generated population. In contrast to the achieved observables, several calibrated parameter distributions differ substantially between methods. The largest location and dispersion shifts occur in the cardiac elastance parameters (*E*_*Hl*_, *E*_*Hr*_) and arterial compliances (*C*_*As*_, *C*_*Ap*_), where the NN exhibits strong positive median shifts and inflated dispersion, indicating systematic re-parameterisation rather than small perturbations about the EGD solution.

**Table 6.** Distributional equivalence metrics between EGD and NN calibrated parameters. For each parameter, the table reports the relative median shift 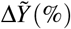, the interquartile range (IQR) ratio, the first-order Wasserstein distance *W*_1_, and the Kolmogorov–Smirnov (KS) statistic comparing the NN and EGD population distributions.

| Variable | $\Delta\tilde{Y}$ (%) | IQR ratio | $W_1$ | KS |
| --- | --- | --- | --- | --- |
| $C_{Vs}$ | -1.80 | 0.984 | 0.935 | 0.052 |
| $R_{ApCp}$ | -3.09 | 0.869 | 0.0196 | 0.110 |
| $R_{CsVs}$ | -1.06 | 1.037 | 0.0155 | 0.040 |
| $R_{AsCs}$ | -5.51 | 0.968 | 0.0467 | 0.080 |
| $E_{Hl}$ | +17.70 | 1.583 | 3.13 | 0.202 |
| $E_{Hr}$ | +13.09 | 1.423 | 0.572 | 0.158 |
| $C_{As}$ | +22.32 | 1.109 | 0.0611 | 0.365 |
| $C_{Ap}$ | +20.37 | 1.208 | 0.176 | 0.188 |
| $V_{0As}$ | -4.68 | 0.995 | 3.23 | 0.128 |
| $V_{0Ap}$ | -27.93 | 0.993 | 2.17 | 0.158 |
| $e_{Hl}$ | +8.32 | 1.209 | 0.0352 | 0.145 |
| $e_{Hr}$ | +11.97 | 1.059 | 0.0298 | 0.142 |
| $C_{Cs}$ | -5.88 | 0.979 | 0.0831 | 0.105 |
| $C_{Vt}$ | +1.16 | 0.944 | 0.0659 | 0.068 |
| $C_{Cp}$ | -0.96 | 0.988 | 0.0221 | 0.057 |
| $C_{Vp}$ | +1.29 | 1.286 | 0.0688 | 0.070 |

*V*0_*Ap*_ shows a large negative median shift and elevated *W*_1_/KS, consistent with the qualitative shift and tail behaviour seen in the parameter distribution plots. These results support the interpretation that the NN often reproduces the output distributions while achieving them via different parameter distribution in specific physiological submodules. Across calibrated parameters, the IQR ratio indicates that dispersion is preserved for many variables (IQR ratio ≈ 1 for *C*_*Vs*_, *R*_*CsVs*_, *R*_*AsCs*_, *V*0_*As*_, *V*0_*Ap*_, *C*_*Cs*_, *C*_*Cp*_), but substantial dispersion inflation is observed in key cardiac and arterial parameters, most notably *E*_*Hl*_ (1.583), *E*_*Hr*_ (1.423), *C*_*Vp*_ (1.286), and *C*_*Ap*_ (1.208), indicating that the NN produces a markedly broader parameter spread than EGD in these submodules.

Figure 5 compares EGD-calibrated and NN-inferred values for a subset of parameters. *E*_*Hl*_ is included because it shows the strongest distribution mismatch between methods in Table 6, *V*_0*Ap*_ because it is associated with the largest errors in 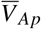, and *C*_*Ap*_ because it plays a complementary role to *V*_0*Ap*_ in the pulmonary arterial pressure–volume relation and therefore captures compensatory reparameterisation within the same submodule.

**Figure 5.**
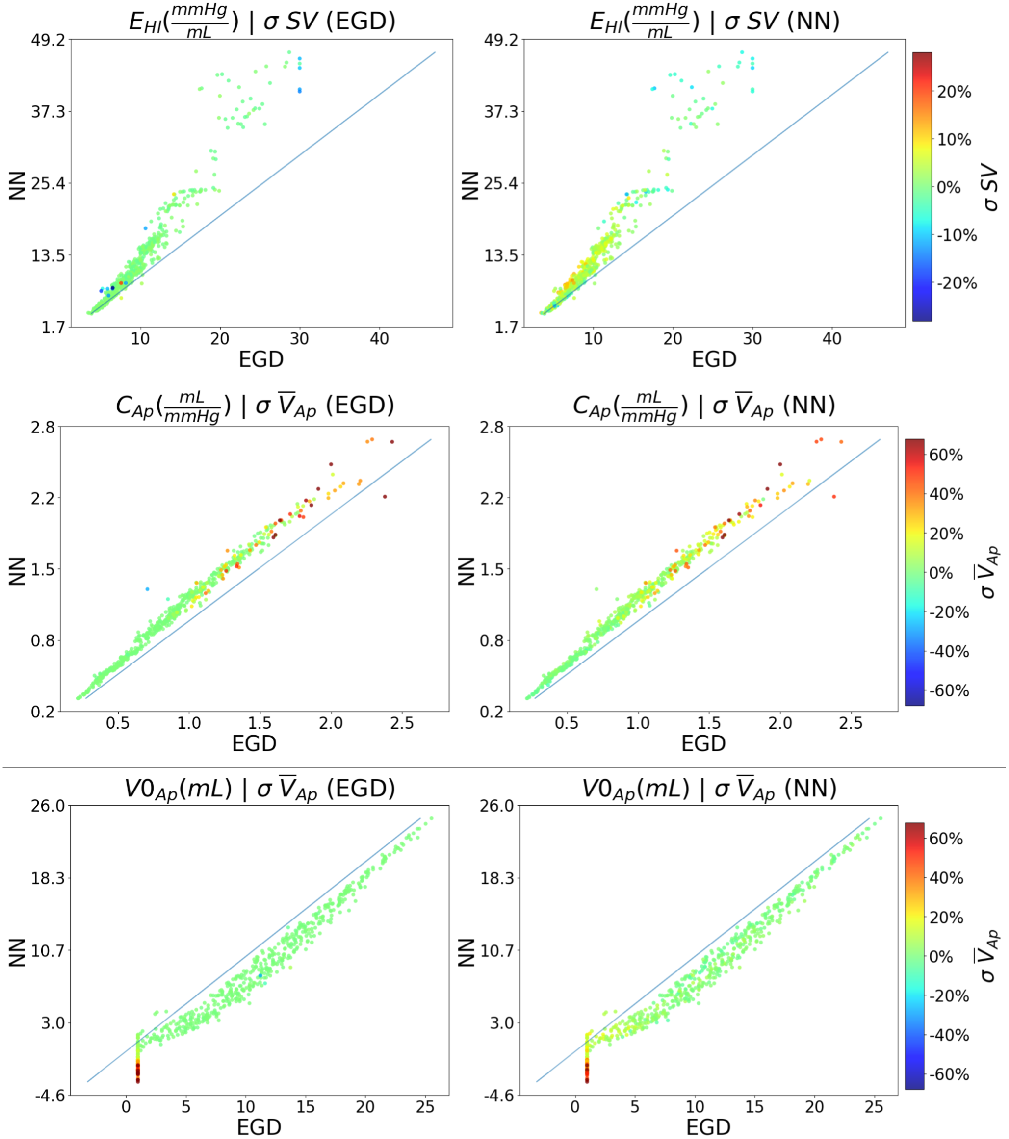
Each row shows an EGD–NN scatter (EGD on the horizontal axis; NN on the vertical axis); the diagonal denotes equality. Top: left-ventricular systolic elastance *E*_*Hl*_ coloured by stroke-volume relative error *σ*_*SV*_ from forward re-simulation. Middle: pulmonary arterial unstressed volume *V*_0*Ap*_ coloured by pulmonary arterial mean-volume relative error 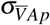. Bottom: pulmonary arterial compliance *CAp* coloured by 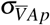. Left panels use errors from EGD-based forward simulations; right panels use errors after re-injecting *θNN* into the mechanistic model, indicating regimes where parameter mismatch coincides with elevated haemodynamic residuals.

Across the population, the NN produced parameter estimates very similar to those obtained with EGD. In most cases, the two methods agreed closely, although the neural network sometimes reached a similar physiological fit by using slightly different combinations of model parameters. the main exception is *E*_*Hl*_, which exhibits a systematic upward shift relative to EGD, indicating that the NN tends to attribute a larger fraction of the required haemodynamic adjustment to heart contractility. In the pulmonary arterial compartment, the NN preferentially infers lower *V*_0*Ap*_ and higher *C*_*Ap*_. The main parameter discrepancies relative to EGD seem to arise in samples that require low pulmonary arterial blood volume 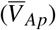 together with high compliance values.

Figure 6 localises where, in haemodynamic target space, the NN produces parameter vectors that are structurally different from the corresponding EGD solutions. Parameter-space disparity is quantified by the cosine distance *D*_cos_ between *θ*_*NN*_ and *θ*_*EGD*_, which captures changes in the relative parameter pattern. Across the virtual population, most samples exhibit very small *D*_cos_, indicating that the NN typically re-produces the EGD parameter-space structure. However, the high-disparity tail (beyond the empirical threshold) is not uniformly distributed across target values: elevated concentrations occur most clearly at elevated systemic pulse pressure *PP*_*As*_, within restricted stroke-volume bands, and at low *TBV* configurations. Point colouring by the NN mean absolute relative error further shows that large *D*_cos_ is more frequently associated with increased aggregate forward error, while a subset of high-disparity cases remains low-error, consistent with some degree of reparameterisation under accurate forward reproduction. Together, these patterns indicate that NN–EGD structural disagreement is target-dependent and concentrated in particular haemodynamic configurations rather than arising randomly across the admissible target space.

**Figure 6.**
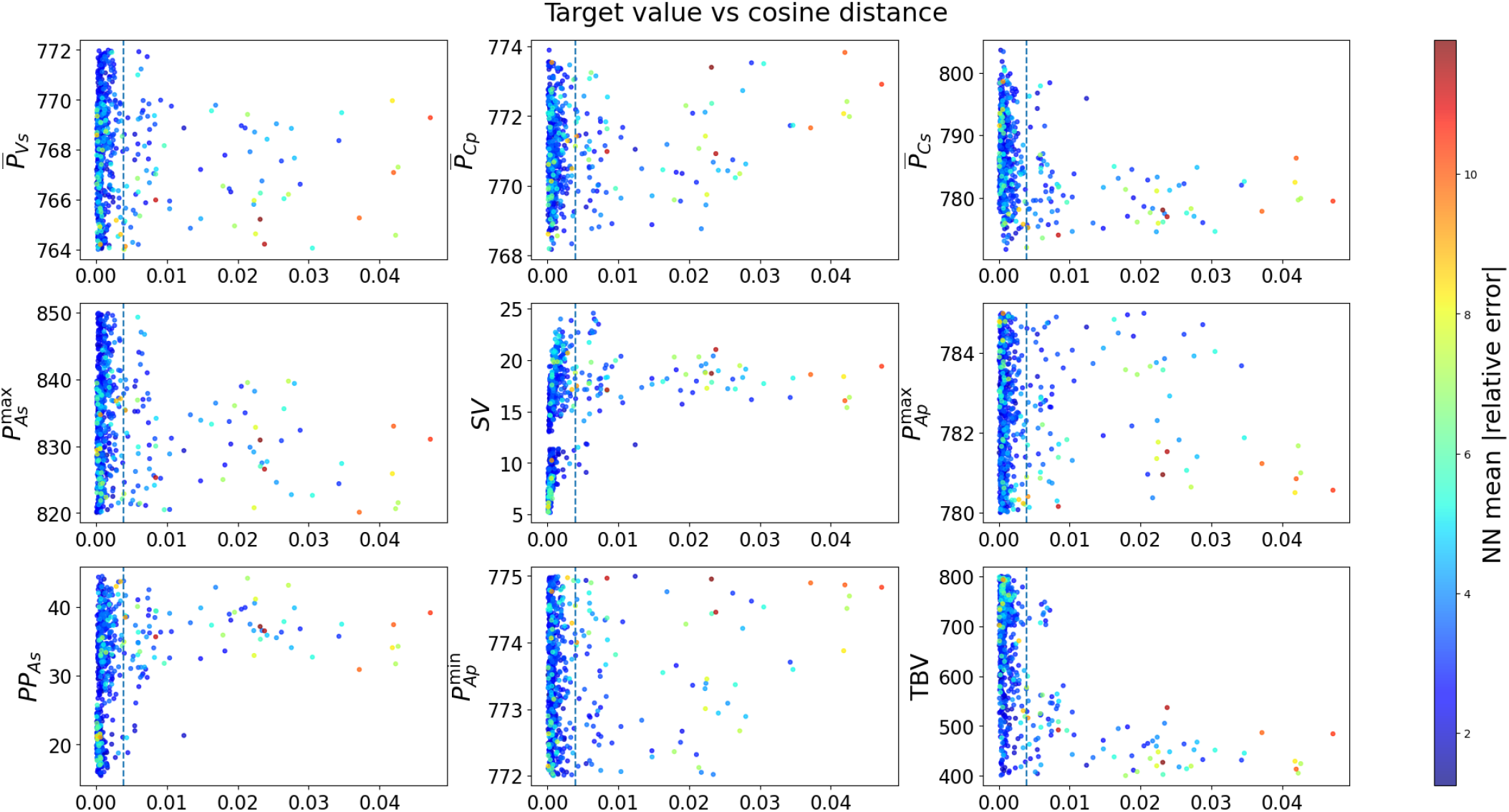
Relationship between NN–EGD parameter-space disparity and haemodynamic regime. For each calibrated sample, the horizontal axis shows the cosine distance between the NN- and EGD-inferred parameter vectors computed over the full calibrated parameter set. Each panel plots *D*_cos_ against a forward-simulated steady-state haemodynamic variable (cycle-averaged pressures, *SV*, pulse pressure *PP*_*As*_, and *TBV*). Points are coloured by the NN mean absolute relative error aggregated across all targets after re-injection of *θ*_*NN*_ into the mechanistic model. The vertical dashed line denotes the top 15% most distant samples.

Figure 7 maps where NN–EGD high-disparity solutions occur within key haemodynamic target spaces. In each row, the binned heatmap reports the fraction of samples exceeding the disparity threshold (cell annotations show bin counts), and the corresponding scatter plot shows the continuous distribution and per-sample disparity magnitude. High-disparity cases concentrate at low-*TBV* configurations with high *SV* and large pulse amplitudes (e.g., high *PP*_*As*_). In the systemic pressure plane, high-disparity points fall predominantly along a distinct high-*PP*_*As*_ branch, consistent with compensatory re-parameterisation confined to specific target regions rather than diffuse mismatch across the population.

**Figure 7.**
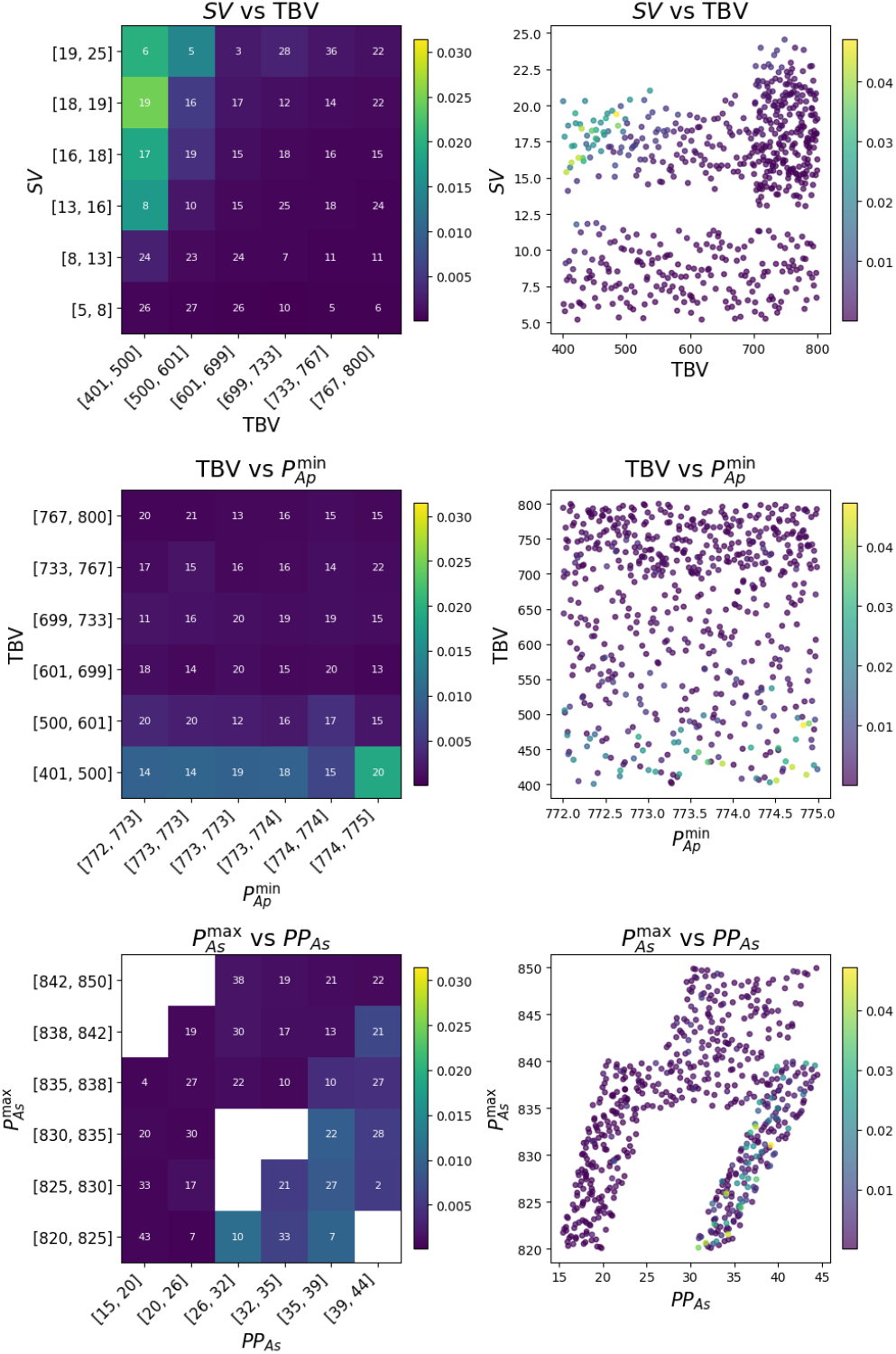
Scatter of prescribed model targets coloured by NN/EGD parameter-space disparity magnitude. For each target pair (row), the right panel shows the continuous target scatter with colour indicating per-sample disparity, while the left panel shows the corresponding binned map of mean disparity in each region of target space (numbers indicate the number of samples per bin). Disparity is concentrated in specific regions rather than uniformly across the population, most notably at low-*TBV*.

## 4 DISCUSSION

This paper presents a neural-network approach for fast calibration of a mechanistic closed-loop cardiovascular model by learning a direct mapping from prescribed haemodynamic target vectors to parameter estimates. The central question is not whether a NN can exactly replicate a simulator-constrained optimiser, but under what conditions a learned inverse can provide reliable parameter proposals while preserving mechanistic plausibility.

To answer this, we use Embedded Gradient Descent (EGD) as a controlled reference on the same model and target sets, and we evaluate agreement by forward re-simulation rather than by parameter matching alone.

The discussion therefore proceeds in three steps. We first interpret the target-tracking behaviour and error structure observed in the scatter/violin summaries (Figure 3 and Table 5), which separate the low-error bulk from a small high-residual subset shared by both methods. We then examine how the remaining differences manifest in parameter space (Table 6 and Figure 4–5), to distinguish local re-parametrisation from method-specific artefacts. Finally, we localise where disagreement concentrates in target space (Figure 6 and Figure 7) to relate failure modes to specific haemodynamic regimes and to motivate practical use of the NN within a mechanistic workflow.

### Validation against EGD

Using EGD as a simulator-constrained reference enables a controlled validation because both methods operate on the same model and prescribed target sets, differing only in how parameters are inferred. Across the full set of observables, EGD yields tighter agreement with the prescribed targets and more concentrated error distributions (Table 5), while the NN reproduces the same overall trends with slightly increased dispersion. The scatter/violin summaries of Figure 3 make this separation explicit; for most targets the predictions of both methods form a dense cloud around the identity line with predominantly small relative errors. The remaining discrepancy in aggregate error metrics is largely driven by a small subset of high-residual outliers, visible as sparse points far from the identity line and as heavy tails in the corresponding violin plots.

Importantly, the same target configurations drive these high-residual cases under both approaches, indicating that the dominant failures reflect feasibility limits of the target/model combination rather than instability of either calibration procedure. This is most evident in the pulmonary arterial volume; Figure 5 shows that, for the high-residual subset, both methods concentrate error at low *V*_0*Ap*_, with EGD solutions saturating at *V*_0*Ap*_ = 0.1 and NN predictions extending into *V*_0*Ap*_ *<* 0 while coinciding with elevated *C*_*Ap*_. Here, the points that lie furthest from the identity line are also those with the largest forward error in 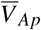, showing that the two methods behave similarly on feasible target sets (low-error samples) but diverge primarily when the targets are incompatible. In those incompatible regimes the NN tends to return parameters that are statistically plausible under its training distribution rather than parameters constrained by mechanistic feasibility, which can yield non-physiological predictions such as *V*_0*Ap*_ *<* 0 (inconsistent with *V*_0*Ap*_ as an unstressed, zero-pressure volume).

### Parameter distribution

The parameter distribution comparison (Table 6 and Figure 4) shows that NN and EGD calibrated populations remain broadly consistent across most of the parameter set. For many parameters the interquartile spread is similar between methods, with IQR ratios close to unity (Table 6), indicating that the bulk of the inferred population structure is preserved. The dominant systematic differences are instead localised to the heart and proximal arterial compartments, where both methods can reproduce similar targets using compensatory adjustments in contractility and arterial load. This is reflected by coherent median shifts and dispersion changes in the heart elastances (*E*_*Hl*_ and *E*_*Hr*_) and in the arterial compliances (*C*_*As*_ and *C*_*Ap*_), consistent with a reparameterisation along a contractility/compliance axis rather than a global redistribution of parameters.

At the same time, the most extreme distribution discrepancies are concentrated within a small subset of cases that also exhibit elevated forward error and large NN–EGD disparity (Figure 3, Figure 6, and Figure 7). In other words, while the NN and EGD populations are closely aligned for the majority of feasible target sets, the high-error tail amplifies the apparent separation for the most sensitive parameter groups. This motivates inspecting the direction of the median shifts and their pairing across *V*_0_, *C*, and *E* parameters to interpret the specific compensatory mechanisms underpinning the observed re-parameterisation.

Inspection of the median shifts suggests two dominant compensation trends. First, the dominant median-shift pattern is a decrease in unstressed volume coupled to an increase in the corresponding compliance, with the effect driven primarily by the pulmonary compartment (notably the large negative shift in *V*_0*Ap*_) while systemic *V*_0_ shifts are comparatively small. This asymmetry is consistent with the observation that the systemic targets are largely feasible under the present model, whereas the pulmonary targets include regimes that are harder to satisfy. Second, the NN tends to favour higher ventricular elastances (equivalently lower ventricular compliances under *C* = 1*/E*) alongside higher arterial compliances. These parameters jointly modulate systolic and pulse pressure generation and transmission, so increased contractility can be partially counterbalanced by increased arterial compliance to attenuate rises in peak and pulse pressure. This compensation is also consistent with the positive median shift in *SV* under NN calibration (Table 5), suggesting that similar target tracking can be obtained while redistributing adjustments across ventricular contractility and arterial compliance degrees of freedom. In addition, Table 6 shows small but consistent shifts towards lower resistances (e.g. in the pulmonary and systemic arterial/venous resistance terms) alongside higher arterial compliances. Although the resistance median shifts are numerically modest, their haemodynamic impact can still be meaningful because pressures and flows depend directly on resistance through the model’s pressure/flow relations, so even small absolute changes can alter the operating point required to match the prescribed targets. Overall, these patterns support an interpretation in terms of local re-parameterisation rather than uniformly degraded calibration.

Finally, the regions of greatest *E*_*Hl*_ discrepancy are also those associated with higher forward errors. As illustrated by Figure 5, a subset of cases with high prescribed *SV* appears infeasible under the available contractility range, with the EGD solution saturating at the upper bound of *E*_*Hl*_. In the NN, the corresponding high-*E*_*Hl*_ predictions coincide with elevated error and systematic underestimation of *SV*, indicating that these points again reflect difficult or incompatible target configurations rather than broad disagreement between the calibration methods.

### Feasibility limits

Because the results indicate both reparameterisation within the feasible regime and a subset of target sets that remain high-residual under both methods, it is important to identify where in target space the NN begins to diverge from the EGD reference and whether this divergence coincides with increased forward error. While distribution-level statistics such as KS and *W*_1_ quantify how much parameter distributions differ overall, they do not indicate which target combinations generate those differences, motivating an explicit localisation analysis in target space. We therefore use the cosine distance *D*_cos_ between *θ*_NN_ and *θ*_EGD_ as a structure-level measure of parameter disagreement that captures changes in the relative parameter pattern, and interpret it jointly with forward re-simulation error to distinguish benign re-parameterisation (high *D*_cos_ but low error) from problematic cases (high *D*_cos_ and high error).

Figure 6 makes this separation explicit: the high-disparity, high-error samples are not spread across the full population, but cluster within narrow bands of the target axes. In particular, the problematic region is characterised by high *SV* at low *TBV*, and by configurations with relatively high *PP*_*As*_ occurring alongside low *SysP*_*As*_, indicating that the dominant failures are driven by specific target combinations rather than diffuse method mismatch. This localisation is consistent with the feasibility-limit interpretation from Figure 3 and Figure 7, where sparse points far from the identity line and heavy-tailed errors occur primarily in restricted regions of target space. Under the current compartment volume distribution assumptions, this target combination cannot be matched while maintaining the model’s closed-loop pressure–volume constraints, so large residuals persist after forward re-simulation.

Finally, Figure 6 shows that these high-error/high-disparity points are dominated by warm shock samples. This is consistent with the target definitions in Table 1, where warm shock combines higher cardiac output with systematically higher systemic pulse pressure targets at the same low systolic pressure range, producing low-diastolic, high-flow configurations that are more likely to approach the structural limits of the present model/target configuration.

### Practical role of the NN

The results support positioning the NN as accelerator rather than a replacement for simulator-constrained calibration, since once trained it replaces iterative simulator-based calibration with a single direct prediction of the parameter vector. A practical workflow is to use the NN to provide an initial parameter proposal, followed by short EGD refinement and forward re-simulation to verify target tracking and flag infeasible target sets. This hybrid approach preserves the primary advantage of the NN, namely rapid initialisation via a single forward pass, while restoring mech-anistic feasibility via EGD refinement and reducing the risk of non-physical parameter configurations. More broadly, the NN is well-suited for screening large target sets, prioritising feasible regions of target space, and reducing reliance on long iterative calibration runs per twin.

### Limitations

Although we trained the NN using broader parameter bounds than those used to generate the evaluation populations, the training distribution still represents only a small subset of the physiologically possible domain. The present work is restricted to the paediatric parameterisation and target ranges considered here and does not attempt to span other domains such as adult physiology or alternative disease spectra. As a result, the NN must be retrained or extended when the model, priors, or target domain changes, whereas EGD can, in principle, be applied to any admissible target set without retraining provided that the forward model remains well-posed.

A second limitation is that the inverse mapping from target summaries to parameters is not guaranteed to be unique. The NN produces a single point estimate for each target vector, yet multiple parameter combinations can yield similar haemodynamic summaries, particularly when the targets are cycle-derived aggregates. In this setting, differences between NN and EGD solutions can reflect alternative but compensatory parameterisations within the feasible regime rather than incorrect calibration.

Physiological constraints are not enforced by construction in the present NN, which can lead to occasional non-physical parameter predictions such as *V*_0*Ap*_ *<* 0. Constraint-aware designs, including bounded parameterisations, positivity-preserving transforms, or loss penalties linked to forward feasibility, may reduce these failures and improve robustness in the challenging regions of target space identified in this study.

Modelling assumptions also play a central role in the observed feasibility limits. A subset of target sets appears incompatible with the current equation system under the present compartment volume distribution used to initialise blood volume allocation. This suggests that, for extreme disease regimes, phenotype-dependent assumptions about blood volume partitioning, or systematic shifts in the *TBV* distribution itself, may be required to expand the feasible target space. A practical advantage of the NN framework is that it makes such sensitivity studies computationally straightforward, for example, by rapidly testing the same prescribed targets under alternative *TBV* distributions and volume-partition priors. In addition, when targets imply strongly time-dependent behaviour, extensions of the forward model that account for neglected dynamics such as blood inertia may be required.

Finally, validation is primarily based on forward resimulation target tracking and distributional comparisons of inferred parameters. These metrics confirm reproduction of the prescribed haemodynamic summaries at a single steady-state operating point, but the cardiovascular system is regulated and time-varying, and the present model does not include autonomic (ANS) dynamics that govern transient adaptation. Consequently, parameter sets that match one snapshot may not yield consistent dynamics or intervention responses when the system is perturbed. Mechanistically anchored validation should therefore extend to standardised in-silico interventions (e.g. volume/afterload challenges or therapy-like perturbations) within a model that includes the relevant regulatory mechanisms, and compare predicted response patterns against direct measurements of observable variables.

## 5 CONCLUSION

This work introduces and evaluates a neural-network approach for fast calibration of a mechanistic closed-loop cardiovascular model, in which prescribed haemodynamic target vectors are mapped directly to parameter estimates. Once trained, the NN returns parameters via a single forward pass and therefore provides orders of magnitude cheaper inference than iterative simulator-constrained calibration that requires repeated ODE integration and parameter updates. Using EGD as a controlled reference and validating by forward re-simulation, we find that NN-based calibration achieves predominantly small target tracking errors for the majority of target sets and preserves much of the population-level parameter structure. The main discrepancies arise in a small subset of target configurations that are difficult or infeasible under the current equation system and modelling assumptions, where both methods exhibit elevated residuals and the NN can produce occasional non-physiological parameter values. Overall, the results support using the NN as a practical accelerator for virtual twin generation and target-space screening, with simulator-based refinement and forward re-simulation retained when tight target tracking and mechanistic plausibility are required.

## CODE AVAILABILITY

The implementation used in this study is available at https://github.com/mtc8608/My-ICU-Twin/tree/NNcalibration

## ACKNOWLEDGMENTS

The authors are grateful for the funding and support of the EPSRC-funded CHIMERA Maths in Healthcare Hub (EP/T017791/1), the Wellcome/EPSRC Centre for Interventional and Surgical Sciences (WEISS) (203145/A/16/Z), the Department of Mechanical Engineering and Department of Mathematics at UCL.

## DECLARATION OF GENERATIVE AI AND AI-ASSISTED TECHNOLOGIES IN THE WRITING PROCESS

During the preparation of this work the author(s) used GPT5 in order to Correct grammar and improve readability. After using this tool/service, the author(s) reviewed and edited the content as needed and take(s) full responsibility for the content of the publication.

## A FULL ERROR COMPARISON

**Figure 8.**
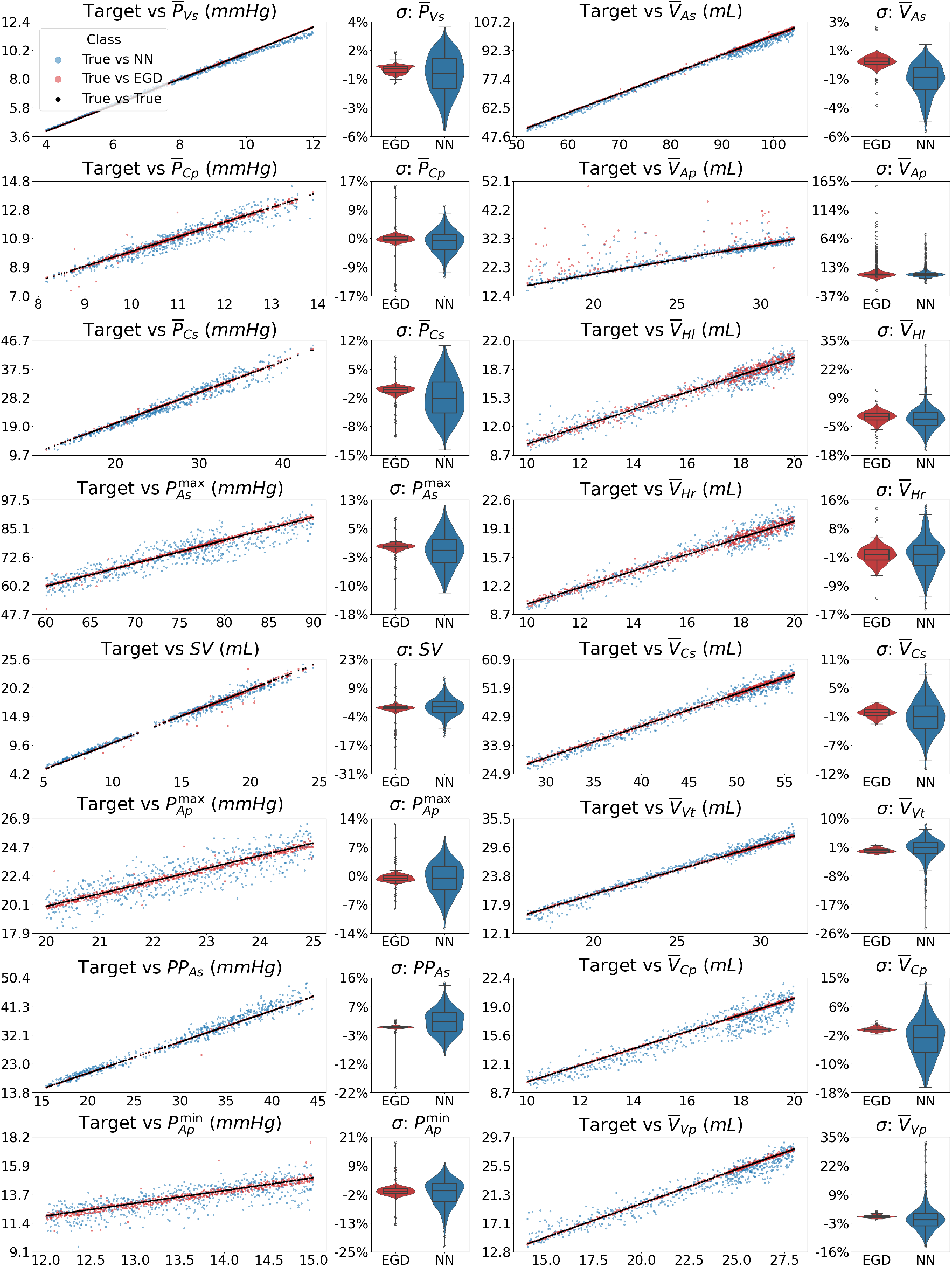
Comparison of calibration fidelity between Embedded Gradient Descent (EGD) and neural-network (NN) inverse calibration across all physiological targets. For each target variable *Y*, the left panel shows the forward-simulated steady-state output versus the prescribed target (identity line), with EGD (red) and NN (blue) overlaid. The right panel shows the corresponding distribution of relative errors *σ*_*Y*_ = (*Y*_model_ − *Y*_target_)*/Y*_target_ across the calibrated population for each method.

## B NOMENCLATURE

**Table 7.** Nomenclature used in the cardiovascular model and calibration framework.

| Symbol | Compartment |
| --- | --- |
| HI | Left ventricle |
| Hr | Right ventricle |
| As | Systemic arteries |
| Ap | Pulmonary arteries |
| Cs | Systemic capillaries |
| Cp | Pulmonary capillaries |
| Vs | Systemic veins |
| Vp | Pulmonary veins |
| Vt | Thoracic veins |
| Symbol | Primary Variable / Parameter |
| $P_*$ | Pressure in compartment * |
| $Q_{* \dagger}$ | Blood flow rate from compartment * to † |
| $V_*$ | Blood volume in compartment * |
| $C_*$ | Compliance of compartment * |
| $R_{* \dagger}$ | Resistance between compartments * and † |
| $V_{0*}$ | Unstressed volume of compartment * |
| $E_{H*}$ | Max (systolic) elastance of heart $H^*$ |
| $e_{H*}$ | Min (diastolic) elastance of heart $H^*$ |
| $\mathcal{E}_{H*}(\tau)$ | Time-varying elastance of heart $H^*$ |
| $R^{in}, R^{out}$ | Generic inlet and outlet resistances |
| $P_{Bias}$ | Pressure offset in capacitor equation |
| $P_{Atm}$ | Atmospheric reference pressure |
| Symbol | Clinical / Derived Quantity |
| $SysP_*$ | Systemic arterial systolic blood pressure |
| $PP$ | Pulse pressure |
| $SV$ | Stroke volume |
| $CO$ | Cardiac output |
| $HR$ | Heart rate |
| $TBV$ | Total blood volume |
| $CVP$ | Central venous pressure |
| Symbol | Auxiliary / Operator Variable |
| $Y(t)$ | Generic scalar physiological/aux. variable |
| $\min(Y)$ | Min of $Y$ over the current cardiac cycle |
| $\max(Y)$ | Max of $Y$ over the current cardiac cycle |
| $\text{avg}(Y)$ | Moving or cycle-average of $Y$ |
| $\text{int}(Y)$ | Integral of $Y$ over the current cardiac cycle |
| $\text{keep}(Y)$ | Value of $Y$ stored from the previous cycle |
| $\mathcal{O}_1(\mathcal{O}_2(Y))$ | Composition of operators acting on $Y$ |
| Symbol | Time Variable |
| $t$ | Absolute simulation time |
| $\tau$ | Intra-cardiac-cycle time (phase variable) |
| $HC$ | Duration of a cardiac cycle |
| $T_{\text{trig}}$ | Trigger time of the next cardiac cycle |
| $T_{\text{avg}}$ | Averaging window duration |
| $\Delta t$ | Numerical integration time step |
| Symbol | Calibration / Controller Variable |
| $Y_{\min}, Y_{\max}$ | Lower and upper bounds of parameter $Y$ |
| $Y_{\text{target}}$ | Target value for variable $Y$ |
| $Y_{\text{current}}$ | Current model value of $Y$ |
| $\sigma_{\text{rel}}$ | Relative error $(Y_{\text{target}} - Y_{\text{current}})/Y_{\text{target}}$ |
| $a$ | Controller gain (scaling factor) |

## C CARDIOVASCULAR MODEL EQUATION OVERVIEW

This appendix provides a minimal specification of the cardiovascular model used in this study. The formulation matches that used in our prior work [16] and is included here solely to define the governing equation system and the cycle-based operators required to compute calibration targets. The model is a closed-loop lumped-parameter network comprising nine compartments: left heart (*Hl*), right heart (*Hr*), systemic arteries (*As*), systemic capillaries (*Cs*), systemic veins (*Vs*), thoracic veins (*Vt*), pulmonary arteries (*Ap*), pulmonary capillaries (*Cp*), and pulmonary veins (*V p*). Each compartment is characterised by a pressure *P*, a volume *V*, and flows to adjacent compartments, following the generic compartment structure illustrated in Figure 9.

**Figure 9.**
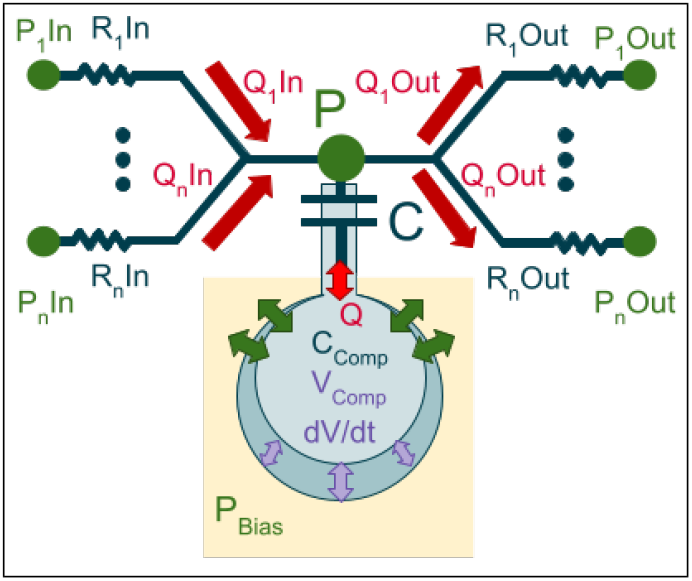
Generic representation of a compartment in the cardiovascular model. Each compartment is connected to its neighbours through a set of resistors, characterised by inlet and outlet resistances *R*_*in*_ and *R*_*out*_, which govern the inflow *Q*_*in*_ and outflow *Q*_*out*_ according to the pressure differences with neighbouring compartments (*P*_*in*_,*P*_*out*_). The central capacitor represents the compartment’s compliance *C*, which stores a volume of fluid and determines the local pressure *P*. Together, these elements define the fundamental pressure/volume/flow relationships used throughout the full model.

### C.1 Generic compartment equations

For a generic compartment with *n* neighbouring connections, the pressure–volume relation is

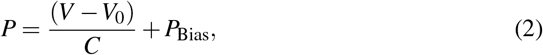

with inflow and outflow defined by linear resistors

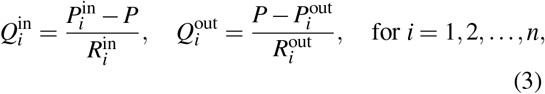

and volume conservation

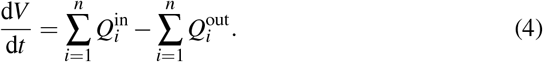

Valvular flows are modelled as diodes

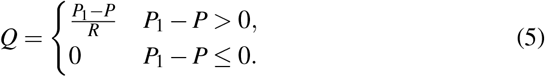

### C.2 Heart elastance and cycle timing

Heart chamber compliances are represented via time-varying elastance *ℰ* (*τ*) = 1*/C*(*τ*), using

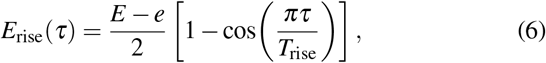

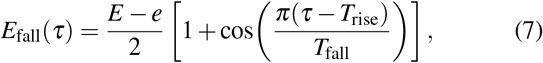

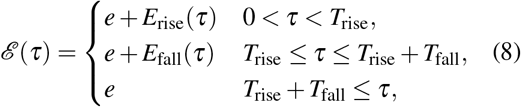

where *E* and *e* are systolic and diastolic elastances and *T*_rise_, *T*_fall_ define contraction and relaxation durations.

The cardiac-cycle duration is

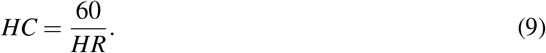

An event-like trigger is implemented with ODE relaxations using solver step Δ*t*:

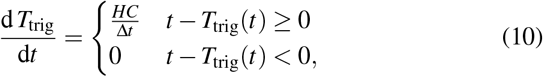

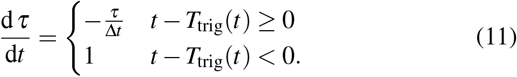

### C.3 Cycle-based operators

Maxima and minima over the current cardiac cycle are computed as

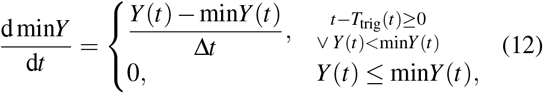

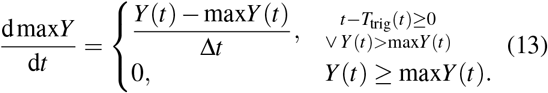

Cycle integrals are computed with reset at each trigger:

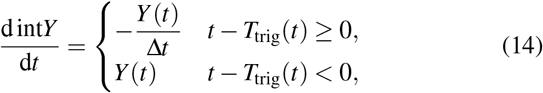

A keeper variable stores the pre-reset value:

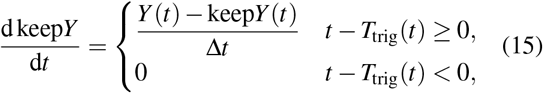

Moving averages over a window *T*_avg_ are computed as

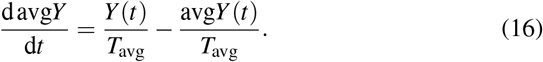

### C.4 Model instantiation

The complete ODE system is obtained by instantiating Eqs. 2– 4 and the inter-compartmental resistive connections for each of the nine compartments, with heart inflow/outflow links modelled using the diode relation Eq. 5. The resulting closed-loop system contains 9 compartmental pressures, 9 compartmental volumes, and 9 inter-compartmental flow variables.

### C.5 Unified model specification

**Table 8.**
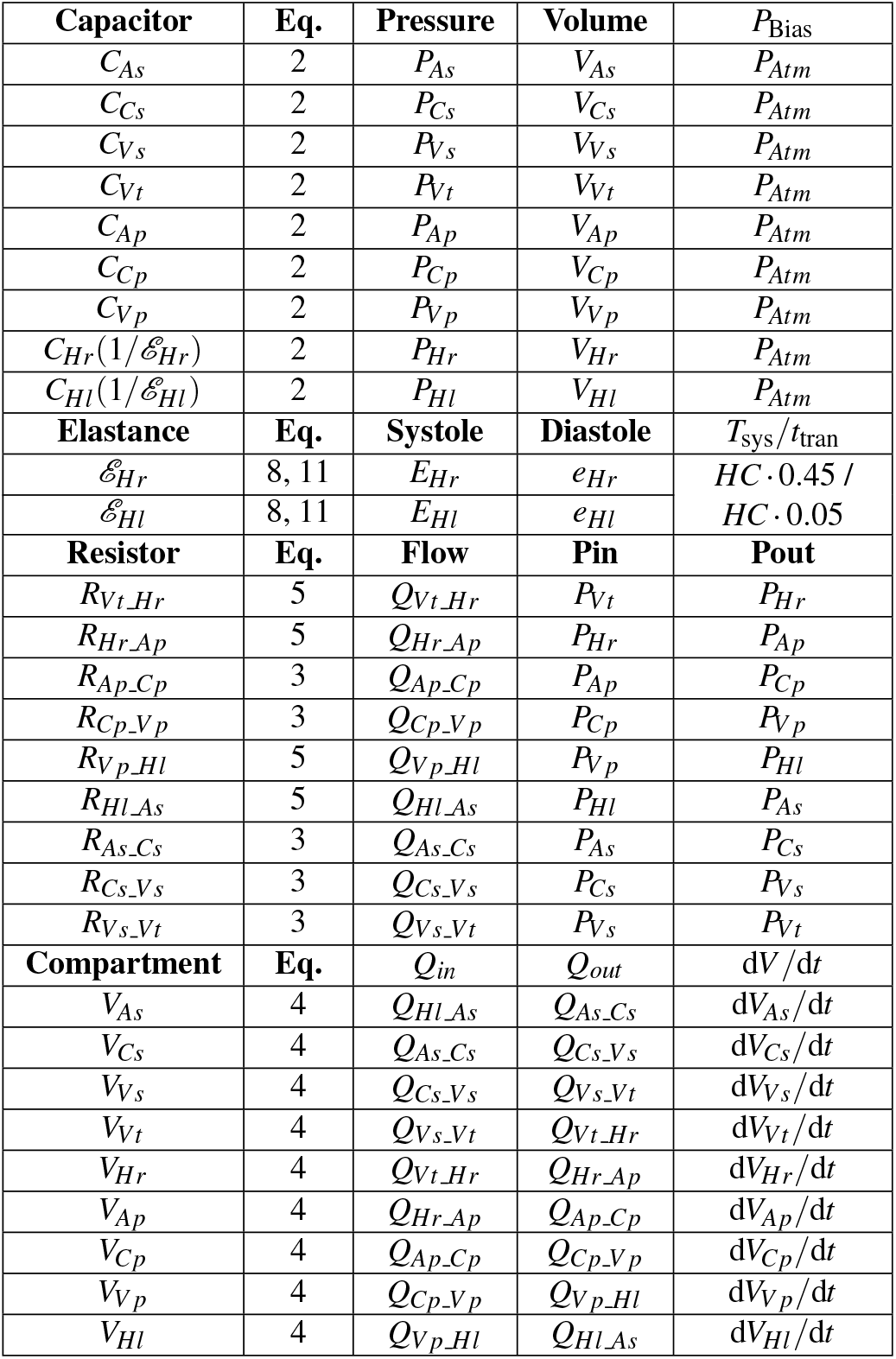
Unified specification of the cardiovascular model. The table defines all compartmental compliance and elastance elements, inter-compartmental resistive connections, and volume balance relationships required to instantiate the full closed-loop system shown in Figure 1.

## Notes

https://github.com/mtc8608/My-ICU-Twin/tree/NNcalibration

### Competing Interest Statement

The authors have declared no competing interest.

